# Co-radiation of *Leptospira* and Tenrecidae (Afrotheria) on Madagascar

**DOI:** 10.1101/2021.12.08.471753

**Authors:** Yann Gomard, Steven M. Goodman, Voahangy Soarimalala, Magali Turpin, Guenaëlle Lenclume, Marion Ah-Vane, Christopher D. Golden, Pablo Tortosa

## Abstract

Leptospirosis is a bacterial zoonosis caused by pathogenic *Leptospira* that are maintained in the kidney lumen of infected animals acting as reservoirs and contaminating the environment via infected urine. The investigation of leptospirosis through a *One Health* framework has been stimulated by notable genetic diversity of pathogenic *Leptospira* together with a high infection prevalence in certain animal reservoirs. Studies of Madagascar’s native mammal fauna have revealed a diversity of *Leptospira* with high levels of host-specificity. Native rodents, tenrecids, and bats shelter several distinct lineages and species of *Leptospira*, some of which have also been detected in acute human cases. Specifically, *Leptospira mayottensis*, first discovered in humans on Mayotte, an island neighboring Madagascar, was subsequently identified in a few species of Malagasy tenrecids, an endemic family of small mammals. Distinct *L. mayottensis* lineages were identified in shrew tenrecs (*Microgale cowani* and *Nesogale dobsoni*) on Madagascar, and later in spiny tenrecs (*Tenrec ecaudatus*) on Mayotte. These findings suggest that *L. mayottensis* (i) has co-radiated with tenrecids on Madagascar, and (ii) has recently emerged in human populations on Mayotte following the introduction of *T. ecaudatus* from Madagascar. Hitherto, *L. mayottensis* has not been detected in spiny tenrecs on Madagascar. In the present study, we broaden the investigation of Malagasy tenrecids and describe the presence of *L. mayottensis* in Malagasy *T. ecaudatus* and *M. thomasi*. These results confirm the hypothesis that *L. mayottensis* was introduced to Mayotte, presumably via *T. ecaudatus*, and provide additional data on the co-radiation of *Leptospira* and Tenrecidae.

## Introduction

Leptospirosis is a zoonotic disease that results annually in around 100 000 human cases and 58 000 deaths [1]. Pathogenic *Leptospira* bacteria are maintained in the lumen of the kidney tubules of animal reservoirs [2], which can chronically shed viable bacteria in their urine and contaminate the environment [3]. Although humans can be infected through direct contact with infected reservoirs, indirect transmission during outdoor activities in the contaminated environment is most frequent [4]. Infection leads to a wide range of symptoms ranging from mild flu-like syndromes to multi-organ failure causing death in 5-10% of the cases.

The genus *Leptospira* is currently composed of more than 60 species including saprophytic and pathogenic species [3, 5–8]. Investigations carried out in different areas of the world through a *One Health* approach have shown distinct transmission chains composed of species or lineages and reservoirs that vary from one environmental setting to another [9–13]. Investigations carried out in the ecosystems of Madagascar and surrounding islands, hereafter referred to as the Malagasy Region, have provided new information on transmission chains on the different islands [14]. Indeed, on La Réunion and in Seychelles, human leptospirosis is mostly caused by *Leptospira* that are broadly distributed and hence likely of introduced origin [11, 12]. By contrast, Madagascar and Mayotte, a French administrated Island belonging to the Comoros archipelago, shelter distinctly more diversified *Leptospira* assemblages, including species and lineages that can be considered endemic [15–17].

Among pathogenic *Leptospira* described and investigated in the Malagasy Region, *L. mayottensis*, the principal focus of the current study, warrants further characterization. These bacteria were first isolated from acute human leptospirosis cases on Mayotte and initially named *L. borgpetersenii* group B [9, 18]. A thorough characterization of serological and genomic features of these isolates led the French Reference Centre on Spirochetes to elevate this bacterium to the rank of a new species, which was named *L. mayottensis* in reference to the geographic origin of the human isolates [19]. A comprehensive investigation of the Malagasy wild mammal fauna allowed identification of *Leptospira* samples imbedded in the genetic clade of *L. mayottensis* and shed by two endemic small mammal species, namely *Microgale cowani* and *Nesogale dobsoni* [20]. These two host species belong to the endemic family Tenrecidae, composed of omnivorous small mammals known to play an important role in *Leptospira* maintenance as reservoirs of two distinct species: *L. borgpetersenii* and *L. mayottensis* [17, 20, 21]. The origin of the Tenrecidae, a monophyletic group, is the result of a single colonization event originating from Africa that took place 30-56 My ago, followed by an extraordinary radiation leading to the currently known nearly 40 extant species or confirmed candidate species [22, 23]. These findings strongly suggest that *L. mayottensis* has co-radiated with tenrecid hosts on Madagascar.

It has been proposed that *L. mayottensis* was introduced to Mayotte from Madagascar [24]. This was supported by an investigation of animal reservoirs on Mayotte identifying *Tenrec ecaudatus*, a spiny tenrec established from Madagascar for human consumption, as the local reservoir of *L. mayottensis*. However, the hypothesis that *T. ecaudatus* sheds *L. mayottensis* currently suffers from a lack of evidence for Malagasy populations of this animal. In the present investigation, we screened *T. ecaudatus* specimens together with other tenrecid species sampled on Madagascar to broaden information on the presence of *L. mayottensis* in these animals, and to test the hypothesis of *L. mayottensis* introduction to Mayotte associated with that of *T. ecaudatus*.

## Materials and Methods

### Biological Sample

All investigated shrew tenrecs were sampled in February 2016 in a forest neighbouring the village of Anjozorobe, in the Central Highlands of Madagascar (see Fig. 1). The samples included 31 specimens belonging to the following nine species: *Microgale taiva* (n=15), *M. thomasi* (n=3), *M. majori* (n=3), *M. parvula* (n=2), *M. soricoides* (n=2), *M. cowani* (n=1), *M. longicaudata* (n=1), *M. fotsifotsy* (n=1) and *Nesogale dobsoni* (n=3). The spiny tenrec samples composed of *T. ecaudatus* included 24 specimens collected in villages adjacent to the Makira Natural Park in the Commune Antsirabe-Sahatany (Maroantsetra District) (Fig. 1), an area with heavy human hunting pressure [25]. All samples in this region were collected from captured animals provided by local hunters to the research team. All specimens were captured, manipulated and euthanized following guidelines accepted by the scientific community for the handling of wild mammals [26] and in strict accordance with permits issued by Malagasy national authorities. All kidney samples from the collected animals from both project areas were immediately stored in ethanol 70% until DNA extraction and molecular analyses.

**Fig. 1:**
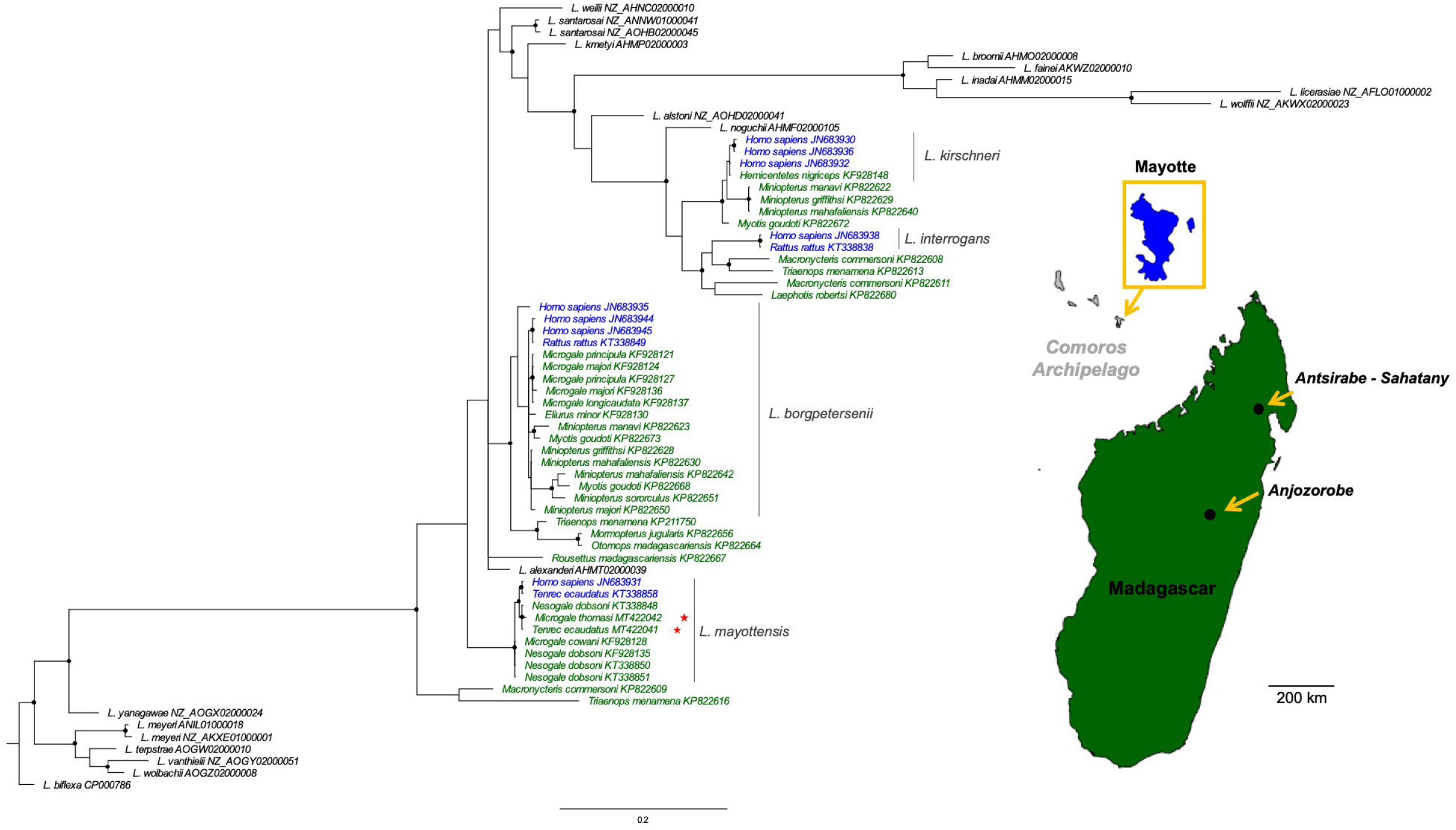
Geographical context and Bayesian phylogenetic tree of *Leptospira* species from Mayotte (blue) and Madagascar (green) based on *secY* gene (482 bp). Sequences in black colour correspond to *Leptosipra* species used as ingroups and outgroup (*L. biflexa*). The accession number is indicated for each sequence. The analysis was conducted under the HKY+I+G substitution model and the tree is midpoint rooted. Black circles at the nodes indicate posterior probabilities superior or equal to 0.90. The red stars indicate new sequences generated in the present study and were obtained from two regions on Madagascar: Anjozorobe and Antsirabe-Sahatany. The map was realised using *worldHires* function in *mapdata* package [42] under the R software version 4.1.1.

### *Leptospira* Detection and Sequencing

For DNA extraction, kidneys were first rinsed with water and subsequently immersed in 2 mL of sterile water overnight. Then a thin transversal slice (approximately 0.5 mm thick) was cut in the central part of the kidney using a sterile scalpel, chopped in small pieces and then immersed into lysis buffer provided in the DNeasy Blood & Tissue Kit (Qiagen, Hilden, Germany) used for DNA extraction. All subsequent extraction steps employed the manufacturer’s instructions. *Leptospira* detection was then carried out on 2 μL of eluted DNA using a probe-specific Real-Time Polymerase Chain Reaction system (RT-PCR) targeting a fragment of the 16S rRNA gene [27]. DNA templates leading to positive RT-PCR results were further subjected to an end-point PCR targeting the *secY* locus as previously described [20]. Amplicons were Sanger sequenced on both strands at GenoScreen (Lille, France) using the same PCR primers. The produced chromatograms were visually edited using Geneious software version 9.0.5 [28].

### Phylogeny

A phylogeny was constructed for the *secY* gene based on the bacterial sequences generated in the present study and previous *secY* sequences from other research in the Malagasy Region [9, 15, 17, 20, 24] (Table S1) and different *Leptospira* species were used as ingroups and outgroups. The best model of sequence evolution was determined with jModelTest v.2.1.4 [29]. Phylogenetic reconstructions were performed with MrBayes v.3.2.3 [30]. The analyses consisted of two independent runs of four incrementally heated Metropolis Coupled Markov Chain Monte Carlo (MCMCMC) starting from a random tree. MCMCMC was run for 2 million generations with trees and associated model parameters sampled every 100 generations. The convergence level of each phylogeny was validated by an average standard deviation of split frequencies inferior to 0.05. The initial 10% of trees for each run were discarded as burn-in and the consensus phylogeny along with posterior probabilities were obtained from the remaining trees. The resulting Bayesian phylogenies were visualized, annotated and rooted to midpoint with FigTree v.1.4.2 [31].

## Results

### Detection of *Leptospira* in Samples and Sequencing

The detection by RT-PCR indicates a global leptospiral infection rate of 7.3% (4/55) with bacteria detected in three out of the nine tested tenrecids species: *M. taiva* (1 positive specimen), *M. thomasi* (2 positive specimens), and *T. ecaudatus* (1 positive specimen). The PCR protocols allowed to obtain leptospiral *secY* sequences from the RT-PCR positive *T. ecaudatus* and from one out of the two RT-PCR positive *M. thomasi*. No *secY* sequence was obtained from the second RT-PCR positive *M. thomasi* or from the RT-PCR positive *M. taiva*.

### Phylogenetic Analysis

We present in Fig. 1 the Bayesian phylogeny obtained from the *secY* gene. Within this phylogeny, the bacterial sequences obtained from *T. ecaudatus* and *M. thomasi* fall in the *L. mayottensis* clade and form a well-supported subclade with a leptospiral sequence obtained from *N. dobsoni*. This subclade is related to one subclade of *L. mayottensis* detected in humans and tenrecs from Mayotte. All previously reported *Leptospira* sequences from *Microgale* and *Nesogale* species fall within two distinct clades: *L. borgpetersenii* (*M. longicaudata, M. principula* and *M. majori*) and *L. mayottensis* (*M. cowani* and *N. dobsoni*). Our results further support this topology with the detection of *L. mayottensis* in *M. thomasi* and Malagasy populations of *T. ecaudatus*.

## Discussion

The Tenrecidae are placental mammals grouped within a monophyletic family endemic to Madagascar and composed of nearly 40 species, including confirmed candidate species [22, 23, 32]. This highly diversified family is currently considered the result from a single colonization event originating from East Africa that took place between 30 and 56 My ago, followed by speciation that resulted in an exceptional adaptive radiation [33, 34]. Some tenrecids exhibit a number of biological features unique among mammals, such as the ability of hibernating without interbout arousal, partial heterothermy or elementary echolocation [32, 35].

The long evolutionary history of the Tenrecidae also makes these mammals suitable for investigating the evolution of host-parasite interactions. Tenrecids host a diversity of Paramyxoviruses, some of which having experienced host switches with introduced murid rodents [36]. Tenrecidae are known to be hosts of two species of pathogenic *Leptospira*, namely *L. borgpetersenii* and *L. mayottensis* [17, 20, 24]. While *L. mayottensis* has been largely identified in tenrecids or acute human cases, a study on Madagascar reported the presence of *L. mayottensis* in introduced *Rattus rattus* but only as co-infections with other *Leptospira* species [37]. The strong host-specificity of *L. mayottensis* towards tenrecids was recently tested through experimental infection in which *L. mayottensis* isolated from *T. ecaudatus* failed to colonize the kidneys of *R. norvegicus* [38]. The present study was carried out to (i) further explore the diversity of *L. mayottensis* sheltered by tenrecids and (ii) confirm a previous hypothesis that proposed *L. mayottensis* arrived on Mayotte with the introduction of *T. ecaudatus* for human consumption.

Analysed samples confirmed tenrecids as being a reservoir of *L. mayottensis* and added *M. thomasi* to the list of animal reservoirs of this pathogenic bacteria. Of particular importance, we report the first characterization of *L. mayottensis* from *T. ecaudatus* on Madagascar. Together with previous data reported on Mayotte [24], the present work supports the introduction of this mammal species being associated with the emergence of a zoonotic human pathogen, *L. mayottensis*, on Mayotte. *Tenrec ecaudatus* has also been introduced to other islands in the Malagasy Region with the purpose of providing bush meat, most notably La Réunion, Mauritius, Mahé (Seychelles) and other islands in the Comoros archipelago, but to our knowledge *L. mayottensis* has been not isolated in these non-native *T. ecaudatus* populations or reported in local human inhabitants.

In conclusion, the data presented herein also strongly support that *L. mayottensis* is an endemic zoonotic pathogen to Madagascar. It has been hypothesized nearly a century ago that the extreme abundance and unbounded dispersal capacities of microorganisms limit endemism, with the exception of some extreme environments, and that biogeographical patterns result from contemporary selective pressures rather than from limited dispersal capacity. This dogma, often referred to as Bass Becking hypothesis - “*everything is everywhere but the environment selects*” [39] - has been increasingly challenged, but microbial biogeography is still in its infancy [40, 41]. The present study supports that host-specificity needs be considered as a driver of microbial endemism: the dispersal capacities of host-specific microbes is indeed limited by that of their hosts. In other words, when considering host-parasite pairs, the dispersal capacities of hosts drive the biogeographical patterns of their associated microorganisms and may, in the case of strong host-parasite specificity, lead to microbial endemism.

## Supporting information

Host families and species, geographical origins, Leptospira species and secY GenBank accession number of Leptospira from Madagascar and Mayotte

## Acknowledgements

We thank the Malagasy authorities (in particular to the former Ministère de l’Environnement, Ecologie et des Forêts, now Ministère de l’Environnement et du Développement Durable) for granting the research permits.

## Statements and Declarations

### Funding

The authors declare that no funds, grants, or other support were received during the preparation of this manuscript.

### Competing interests

The authors declare that they have no competing interest.

### Author Contributions

Pablo Tortosa, Steven M. Goodman and Christopher D. Golden conceived the study. Steven M. Goodman and Christopher D. Golden and Voahangy Soarimalala collected the samples on the field. Magali Turpin, Guenaëlle Lenclume, Marion Ah-Vane and Yann Gomard performed laboratory manipulations. Yann Gomard performed the analysis. Pablo Tortosa, Yann Gomard, Steven M. Goodman and Christopher D. Golden drafted the first version of the manuscript. All authors contributed to the final version of the manuscript.

### Data availability

The produced sequences were deposited in GenBank under the accession numbers MT442041 and MT442042.

### Ethics approval

Biological sampling permits were obtained from the Ministry of Environment and Forests and registered under the following references 20/16/MEEMF/SG/DGF/DAPT/SCBT.Re and 85/18/MEEF/SG/DGF/DSAP/SCB.Re.

## Supplementary materials

**Table S1** Details on *Leptospira secY* sequences from Mayotte and Madagascar used in the present study: host families and species, geographical origins, *Leptospira* species and GenBank accession number.

The asterisks indicate sequences generated in the present study.

## Notes

### Competing Interest Statement

The authors have declared no competing interest.

### Summary of Updates

Leptospirosis is a bacterial zoonosis caused by pathogenic Leptospira that are maintained in the kidney lumen of infected animals acting as reservoirs and contaminating the environment via infected urine. The investigation of leptospirosis through a One Health framework has been stimulated by notable genetic diversity of pathogenic Leptospira together with a high infection prevalence in certain animal reservoirs. Studies of Madagascar native mammal fauna have revealed a diversity of Leptospira with high levels of host-specificity. Native rodents, tenrecids, and bats shelter several distinct lineages and species of Leptospira, some of which have also been detected in acute human cases. Specifically, Leptospira mayottensis, first discovered in humans on Mayotte, an island neighboring Madagascar, was subsequently identified in a few species of Malagasy tenrecids, an endemic family of small mammals. Distinct L. mayottensis lineages were identified in shrew tenrecs (Microgale cowani and Nesogale dobsoni) on Madagascar, and later in spiny tenrecs (Tenrec ecaudatus) on Mayotte. These findings suggest that L. mayottensis (i) has co-radiated with tenrecids on Madagascar, and (ii) has recently emerged in human populations on Mayotte following the introduction of T. ecaudatus from Madagascar. Hitherto, L. mayottensis has not been detected in spiny tenrecs on Madagascar. In the present study, we broaden the investigation of Malagasy tenrecids and describe the presence of L. mayottensis in Malagasy T. ecaudatus and M. thomasi. These results confirm the hypothesis that L. mayottensis was introduced to Mayotte, presumably via T. ecaudatus, and provide additional data on the co-radiation of Leptospira and Tenrecidae.

## References

1. Costa F, Hagan JE, Calcagno J, et al (2015) Global morbidity and mortality of leptospirosis: a systematic review. PLoS Negl Trop Dis 9:e0003898. https://doi.org/10.1371/journal.pntd.0003898

2. Ristow P, Bourhy P, Kerneis S, et al (2008) Biofilm formation by saprophytic and pathogenic leptospires. Microbiol 154:1309–1317. https://doi.org/10.1099/mic.0.2007/014746-0

3. Adler B (2015) Leptospira and Leptospirosis. Springer-Verlag, Berlin Heidelberg

4. Ko AI, Goarant C, Picardeau M (2009) *Leptospira*: the dawn of the molecular genetics era for an emerging zoonotic pathogen. Nat Rev Microbiol 7:736–747. https://doi.org/10.1038/nrmicro2208

5. Masuzawa T, Sakakibara K, Saito M, et al (2018) Characterization of *Leptospira* species isolated from soil collected in Japan. Microbiolo and Immunol 62:55–59. https://doi.org/10.1111/1348-0421.12551

6. Thibeaux R, Iraola G, Ferrés I, et al (2018) Deciphering the unexplored *Leptospira* diversity from soils uncovers genomic evolution to virulence. Microb Genom 4:e000144. https://doi.org/10.1099/mgen.0.000144

7. Thibeaux R, Girault D, Bierque E, et al (2018) Biodiversity of environmental *Leptospira*: improving identification and revisiting the diagnosis. Front Microbiol 9:816. https://doi.org/10.3389/fmicb.2018.00816

8. Vincent AT, Schiettekatte O, Goarant C, et al (2019) Revisiting the taxonomy and evolution of pathogenicity of the genus *Leptospira* through the prism of genomics. PLoS Negl Trop Dis 13:e0007270. https://doi.org/10.1371/journal.pntd.0007270

9. Bourhy P, Collet L, Lernout T, et al (2012) Human *Leptospira* isolates circulating in Mayotte (Indian Ocean) have unique serological and molecular features. J Clin Microbiol 50:307–311. https://doi.org/10.1128/JCM.05931-11

10. Cosson J-F, Picardeau M, Mielcarek M, et al (2014) Epidemiology of *Leptospira* transmitted by rodents in southeast Asia. PLoS Negl Trop Dis 8:e2902. https://doi.org/10.1371/journal.pntd.0002902

11. Guernier V, Lagadec E, Cordonin C, et al (2016) Human leptospirosis on Reunion Island, Indian Ocean: are rodents the (only) ones to blame? PLoS Negl Trop Dis 10:e0004733. https://doi.org/10.1371/journal.pntd.0004733

12. Biscornet L, Dellagi K, Pagès F, et al (2017) Human leptospirosis in Seychelles: a prospective study confirms the heavy burden of the disease but suggests that rats are not the main reservoir. PLoS Negl Trop Dis 11:e0005831. https://doi.org/10.1371/journal.pntd.0005831

13. Guernier V, Richard V, Nhan T, et al (2017) *Leptospira* diversity in animals and humans in Tahiti, French Polynesia. PLoS Negl Trop Dis 11:e0005676. https://doi.org/10.1371/journal.pntd.0005676

14. Gomard Y, Dellagi KM, Goodman S, et al (2021) Tracking animal reservoirs of pathogenic *Leptospira*: the right test for the right claim. Trop Med Infect Dis 6:205. https://doi.org/10.3390/tropicalmed6040205

15. Gomard Y, Dietrich M, Wieseke N, et al (2016) Malagasy bats shelter a considerable genetic diversity of pathogenic *Leptospira* suggesting notable host-specificity patterns. FEMS Microbiol Ecol. https://doi.org/10.1093/femsec/fiw037

16. Tortosa P, Dellagi K, Mavingui P (2017) Leptospiroses on the French islands of the Indian Ocean. Bulletin Epidemiologique Hebdomadaire - BEH 8–9

17. Dietrich M, Gomard Y, Lagadec E, et al (2018) Biogeography of *Leptospira* in wild animal communities inhabiting the insular ecosystem of the western Indian Ocean islands and neighboring Africa. Emerg Microbes Infect 7:57. https://doi.org/10.1038/s41426-018-0059-4

18. Bourhy P, Collet L, Clément S, et al (2010) Isolation and characterization of new *Leptospira* genotypes from patients in Mayotte (Indian Ocean). PLoS Negl Trop Dis 4:e724. https://doi.org/10.1371/journal.pntd.0000724

19. Bourhy P, Collet L, Brisse S, Picardeau M (2014) *Leptospira mayottensis* sp. nov., a pathogenic species of the genus *Leptospira* isolated from humans. Int J Syst Evol Microbiol 64:4061–4067. https://doi.org/10.1099/ijs.0.066597-0

20. Dietrich M, Wilkinson DA, Soarimalala V, et al (2014) Diversification of an emerging pathogen in a biodiversity hotspot: *Leptospira* in endemic small mammals of Madagascar. Mol Ecol 23:2783–2796. https://doi.org/10.1111/mec.12777

21. Dietrich M, Gomard Y, Cordonin C, Tortosa P (In press) Leptospira bacteria in Madagascar. In: The new natural history of Madagascar, S.M Goodman (ed), Princeton University Press. Princeton

22. Everson KM, Soarimalala V, Goodman SM, Olson LE (2016) Multiple loci and complete taxonomic sampling resolve the phylogeny and biogeographic history of Tenrecs (Mammalia: Tenrecidae) and reveal higher speciation rates in Madagascar’s humid forests. Syst Biol 65:890–909. https://doi.org/10.1093/sysbio/syw034

23. Goodman SM, Soarimalala V, Olson LE (2018) Systématique des tenrecs endémiques malgaches (famille des Tenrecidae) / Systematics of endemic Malagasy tenrecs (family Tenrecidae). In: Les aires protégées terrestres de Madagascar: leur histoire, description, et biote. / The terrestrial protected areas of Madagascar: Their history, description, and biota, S.M Goodman, M. J., and S. Wohlhauser (eds). Association Vahatra, pp 363–372

24. Lagadec E, Gomard Y, Minter GL, et al (2016) Identification of *Tenrec ecaudatus,* a wild mammal introduced to Mayotte Island, as a reservoir of the newly identified human pathogenic *Leptospira mayottensis*. PLoS Negl Trop Dis 10:e0004933. https://doi.org/10.1371/journal.pntd.0004933

25. Annapragada A, Brook CE, Luskin MS, et al (2021) Evaluation of tenrec population viability and potential sustainable management under hunting pressure in northeastern Madagascar. Anim Conserv 24:1059–1070. https://doi.org/10.1111/acv.12714

26. Sikes R, Gannon W, The Animal Care and Use Committee of the American Society of Mammalogists (2011) Guidelines of the American Society of Mammalogists for the use of wild mammals research. J Mammal 92:235–253. https://doi.org/10.1644/10-MAMM-F-355.1

27. Smythe LD, Smith IL, Smith GA, et al (2002) A quantitative PCR (TaqMan) assay for pathogenic *Leptospira* spp. BMC Infect Dis 2:13. https://doi.org/10.1186/1471-2334-2-13

28. Kearse M, Moir R, Wilson A, et al (2012) Geneious Basic: an integrated and extendable desktop software platform for the organization and analysis of sequence data. Bioinformatics 28:1647–1649. https://doi.org/10.1093/bioinformatics/bts199

29. Darriba D, Taboada GL, Doallo R, Posada D (2012) jModelTest 2: more models, new heuristics and parallel computing. Nat Methods 9:772–772. https://doi.org/10.1038/nmeth.2109

30. Ronquist F, Teslenko M, van der Mark P, et al (2012) MrBayes 3.2: efficient Bayesian phylogenetic inference and model choice across a large model space. Syst Biol 61:539–542. https://doi.org/10.1093/sysbio/sys029

31. Rambaut A (2014) FigTree 1.4.2 software. Institute of Evolutionary Biology, University of Edinburgh

32. Olson LE (2013) Tenrecs. Curr Biol 23:R5–R8. https://doi.org/10.1016/j.cub.2012.11.015

33. Emerson BC (2002) Evolution on oceanic islands: molecular phylogenetic approaches to understanding pattern and process. Mol Ecol 11:951–966. https://doi.org/10.1046/j.1365-294x.2002.01507.x

34. Krause DW (2010) Washed up in Madagascar. Nature 463:613–614. https://doi.org/10.1038/463613a

35. Lovegrove BG, Lobban KD, Levesque DL (2014) Mammal survival at the Cretaceous–Palaeogene boundary: metabolic homeostasis in prolonged tropical hibernation in tenrecs. Proc Biol Sci 281:20141304. https://doi.org/10.1098/rspb.2014.1304

36. Wilkinson DA, Mélade J, Dietrich M, et al (2014) Highly diverse Morbillivirus-related paramyxoviruses in wild fauna of the southwestern Indian Ocean islands: evidence of exchange between introduced and endemic small mammals. J Virol JVI.01211-14. https://doi.org/10.1128/JVI.01211-14

37. Moseley M, Rahelinirina S, Rajerison M, et al (2018) Mixed *Leptospira* Infections in a diverse reservoir host community, Madagascar, 2013–2015. Emerg Infect Dis 24:1138–1140. https://doi.org/10.3201/eid2406.180035

38. Cordonin C, Turpin M, Bringart M, et al (2020) Pathogenic *Leptospira* and their animal reservoirs: testing host specificity through experimental infection. Sci Rep 10:7239. https://doi.org/10.1038/s41598-020-64172-4

39. Baas Becking L (1934) Geobiologie of Inleiding Tot de Milieukunde, W. P. Van Stockum&Zoon. The Hague, Netherlands

40. de Wit R, Bouvier T (2006) ‘Everything is everywhere, but, the environment selects’; what did Baas Becking and Beijerinck really say? Environ Microbiol 8:755–758. https://doi.org/10.1111/j.1462-2920.2006.01017.x

41. Fierer N (2008) Microbial biogeography: patterns in microbial diversity across space and time. In: Zengler K (ed) Accessing uncultivated microorganisms: from the environment to organisms and genomes and back. Amer Soc Microbiology, Washington, pp 95–115

42. Becker RA, Wilks AR (2018) R version by Ray Brownrigg. mapdata: Extra Map Databases (version 2.3.0)

